# Heterogeneity and Ontogeny of Mouse Thymic Macrophages Reveal a Requirement for *Csf1r*-Expressing Myeloid Cells During Early T Cell Development

**DOI:** 10.1101/2025.08.27.672681

**Authors:** Helen Wang, Vinothkumar Rajan, Anthony Wong, Slava Epelman, Juan Carlos Zúñiga-Pflücker

## Abstract

Thymic macrophages (TMs) maintain tissue homeostasis by clearing the large numbers of apoptotic cells generated during T cell development, but how TM heterogeneity relates to their developmental origin and role in thymocyte maturation remains incompletely understood. Using complementary flow-cytometric, single-cell transcriptomic, and genetic approaches, we resolved two major TM populations corresponding to TIMD4^+^ cortical and CX3CR1^+^ medullary/cortico-medullary macrophages. TIMD4^+^ VCAM1^+^ TMs displayed a prominent efferocytosis and apoptotic-cell-clearance program, whereas TIMD4^−^ VCAM1^+^ TMs were enriched for antigen-presentation and interferon-response pathways. Fate mapping revealed unequal progenitor contributions to these populations, and CCR2 deficiency selectively reduced TIMD4^−^ VCAM1^+^ TMs and thymic monocytes, supporting ongoing input from circulating precursors. Exploratory pseudotime analysis further identified a transcriptional continuum from *Ly6c2*^+^ *Ccr2*^+^ monocytes toward macrophage states. Using MaFIA fetal thymic organ cultures, AP20187-mediated depletion of *Csf1r-*expressing myeloid cells reduced CD4^+^CD8^+^ thymocyte differentiation and produced a coordinated accumulation of DN3 cells, loss of DN4 cells, and reduction in CD27 expression. These convergent changes identify the DN3-to-DN4 transition as a developmental stage that requires an intact *Csf1r*-expressing myeloid compartment, and establish a functional connection between the thymic myeloid niche and early αβ T cell development. Together, our study refines the phenotypic and developmental organization of mouse TMs and reveals a previously under-appreciated requirement for myeloid-cell support during progression through the β-selection checkpoint.

## INTRODUCTION

The thymus is the primary lymphoid organ dedicated to the generation of a diverse and self-tolerant repertoire of T lymphocytes, a process fundamental to adaptive immunity (1). This intricate T cell developmental program unfolds within a highly organized microenvironment composed of thymic epithelial cells (TECs), endothelial cells, and various hematopoietic populations, which collectively provide the essential signals for thymocyte differentiation, proliferation, and selection (2–4). The developmental journey of a T cell begins when bone marrow-derived thymic seeding progenitors (TSPs) enter the thymus at the cortico-medullary junction (CMJ) and commit to the T cell lineage, initiating a multi-stage differentiation cascade (5–7).

This journey is punctuated by a series of stringent checkpoints that assess the integrity and functionality of the nascent T cell receptor (TCR) repertoire (8,9). The earliest stages occur in the thymic cortex, where progenitors, lacking expression of CD4 and CD8, termed double-negative (DN) thymocytes (6), start to differentiate, which is characterized by the expression of CD44 and CD25 (3,4). Progression from the DN1 (CD44^+^CD25^−^) to the DN2 (CD44^+^CD25^+^) and DN3 (CD44^−^CD25^+^) stages is marked by the initiation of V(D)J recombination at the *Tcrb*, *Tcrg*, and *Tcrd* gene loci (1). The DN3 stage represents the first critical checkpoint in αβ T cell development, known as β-selection (12,13). Here, thymocytes that have successfully rearranged a functional TCRβ chain pair it with the invariant pre-TCRα chain (pTα) to form the pre-TCR complex. Successful signaling through this complex provides vital survival, proliferation, and differentiation cues, allowing the cell to progress to the DN4 stage and subsequently to the CD4^+^CD8^+^ double-positive (DP) stage (14,15). Thymocytes failing to generate a functional pre-TCR are eliminated via apoptosis, ensuring that only cells with a viable TCRβ chain continue their developmental progression (14).

Following β-selection, DP thymocytes undergo further TCRα chain rearrangement and are subjected to two additional selection events. Positive selection ensures that the fully assembled αβ TCR can recognize self-peptides presented by major histocompatibility complex (MHC) molecules on cortical (c)TECs, a process essential for TCR MHC-restriction (16–18). Subsequently, negative selection in the medulla eliminates thymocytes whose TCRs bind with high affinity to self-peptides presented by medullary (m)TECs and dendritic cells (DCs), thereby establishing central tolerance (19,20). These selection processes are notably stringent, resulting in the apoptotic death of over 95% of all developing thymocytes (21).

Integral to this journey and processes are thymic macrophages (TMs), which have long been recognized for their critical homeostatic function. Residing in both the cortex and medulla, TMs are exceptionally efficient phagocytes, responsible for the swift and silent clearance of the millions of apoptotic thymocytes generated daily (22–25). This process, known as efferocytosis, is vital for preventing the release of potentially immunogenic and inflammatory intracellular contents and maintaining the integrity of the thymic architecture (26). However, this view of TMs as mere “housekeepers” is rapidly evolving. Recent advances, particularly single-cell RNA sequencing (scRNA-seq), have revealed significant heterogeneity within the TM compartment, suggesting functional specialization (27). Two principal subsets have been identified in the adult mouse thymus: cortical TIMD4+ macrophages, which have potent efferocytosis function, and CX3CR1+ macrophages, which are enriched at the CMJ and express genes associated with antigen presentation, suggesting at a potential role in negative selection (27).

Recent work by Zhou et al. established that adult mouse TMs comprise embryonically derived TIMD4+ macrophages localized predominantly in the cortex and adult hematopoietic stem cell-derived CX3CR1+ macrophages enriched in the medulla and cortico-medullary junction (27). These findings provided an important framework for TM heterogeneity and ontogeny. However, the surface-marker relationships that permit prospective resolution of these populations, the requirements that maintain them, and their contribution to early thymocyte development remain incompletely defined. In particular, β-selection is commonly viewed as a thymocyte-intrinsic checkpoint, even though DN3-to-DN4 progression occurs within a multicellular thymic microenvironment. This raises the possibility that resident myeloid cells provide supportive functions that help developing thymocytes successfully navigate this transition.

Here, we combined high-parameter flow cytometry, single-cell RNA sequencing, genetic fate mapping, CCR2 and SpiC deficiency, and fetal thymic organ culture (FTOC) to define the phenotypic organization, developmental contributions, and functional significance of the thymic myeloid compartment. Our results identify VCAM1 as a useful component of a prospective gating strategy for the two major TM populations, demonstrate CCR2 dependence of the TIMD4-VCAM1+ population, and reveal that depletion of *Csf1r*-expressing myeloid cells compromises thymocyte progression across the DN3-to-DN4 checkpoint.

## MATERIALS AND METHODS

### Animals

Macrophage Fas-Induced Apoptosis (MaFIA), *MafB*-mCherry-Cre, *Spic*^−/−^, and C57BL/6J mice were purchased from the Jackson Laboratory. The *Flt3-*Cre × *Rosa*-mTmG mice and *Flt3-*Cre × *Rosa*-mTmG mice in the *Ccr2^−/-^* background were kindly provided by Dr. Slava Epelman (University Health Network, University of Toronto) (28). All animals were bred and maintained in specific pathogen-free conditions at the Preclinical Research Centre of the Sunnybrook Research Institute (SRI). All animal procedures were approved by the Sunnybrook Health Sciences Centre Animal Care Committee (Toronto, Ontario, Canada).

### Timed pregnancies and embryonic thymus analysis

Timed pregnancies were set up by housing male mice (8–24 weeks of age) in separate cages and mating them with female mice (8–24 weeks of age) overnight. The following morning, females were separated from the males, and successful matings were identified by the presence of a vaginal plug. The gestational age of the embryos was designated as embryonic day 0.5 (E0.5) on the day the plug was observed. Thymuses were harvested from embryos at E15 and E17, as well as from postnatal mice at 1 day, 1 week, and 3 weeks of age, in addition to adult (6-8 weeks) mice. Following collection, thymuses were subjected to enzymatic digestion to dissociate the tissue into single-cell suspensions. The resulting cells were stained for flow cytometric analysis.

### Intravenous CD45 staining

A CD45-Phycoerythrin (PE) antibody (30-F11; BioLegend) was diluted in phosphate buffered saline (PBS) to a final concentration of 1.5 µg/150 µL. The antibody solution was administered via tail vein injection using a 1 mL insulin syringe. Three minutes after the injection, mice were euthanized, and thymuses were harvested. The collected tissue was then processed and stained for flow cytometric analysis.

### Cell isolation from thymus

Thymocytes were harvested and subjected to enzymatic digestion using the Spleen Dissociation Kit (Miltenyi Biotec) and the gentleMACS™ Dissociator (Miltenyi Biotec) according to the manufacturer’s instructions. The resulting cell suspensions were filtered through a 70 µm cell strainer to remove debris, resuspended in PBS, and kept on ice until further processing.

### Flow cytometry

Single-cell suspensions (1 × 10^7^ cells) were prepared from the thymus and stained in a 96-well conical bottom plate (ThermoFisher Scientific). The staining cocktail included the fixable viability dye eFluor 450 (eBioscience), diluted 1:800 in PBS, anti-CD16/32 to block Fc receptors, and the desired fluorochrome-conjugated antibodies targeting macrophages and T cells (see **Supplemental Tables 1 and 2** for details). Cells were initially stained with eFluor 450 in PBS for 15 minutes on ice. After washing with fluorescence-activated cell sorting (FACS) buffer, anti-CD64, anti-CCR2, and anti-VCAM1 antibodies were added to the wells, and cells were stained for an additional 15 minutes on ice. To block non-specific binding, Fc-block antibody was added to the cells 15 minutes prior to the addition of the remaining antibody cocktail, which was incubated with the cells for 30 minutes on ice.

Following staining, cells were washed and resuspended in FACS buffer for flow cytometric acquisition using the FACSymphony™ A5 Cell Analyzer (BD Biosciences). Data were analyzed with FlowJo software (TreeStar). The complete list of antibodies and their corresponding clones is provided in **Supplemental Tables 1 and 2**.

### Cell sorting

Thymuses were harvested from 6-week-old C57BL/6 mice and subjected to enzymatic digestion using the Spleen Dissociation Kit (Miltenyi Biotec) and the gentleMACS™ Dissociator (Miltenyi Biotec) to generate single-cell suspensions. After cell isolation, cells were counted and stained with allophycocyanin (APC)-conjugated anti-mouse CD64 antibody in FACS buffer for 20 minutes on ice (see Supplemental Table 1 for antibody details). Following a wash with FACS buffer, cells were stained with anti-APC MicroBeads (Miltenyi Biotec) for 20 minutes on ice. The cells were then washed again with FACS buffer and loaded onto LS columns (Miltenyi Biotec) for positive selection of CD64^+^ cells. After selection, the cells were washed and further stained with antibodies against F4/80, TIMD4, Ly6C, and CD45 for 30 minutes on ice (see Supplemental Table 1 for antibody details). The stained cells were sorted using the FACSAria™ Fusion Flow Cytometer (BD Biosciences). CD64^+^ sorted cells were collected in 1.5 mL Eppendorf tubes containing 300 µL of FBS, counted, and resuspended in PBS supplemented with 0.04% BSA. Cells were then encapsulated, and library preparation was performed using the Chromium Single Cell 3’ Reagent Kit (10X Genomics). Gene expression libraries were sequenced on a NovaSeq 6000 platform (Illumina) at Azenta Life Sciences. In total, 3,214 and 2,815 cells were sequenced from two independent experiments.

### Fetal thymus organ culture (FTOC) and TM depletion in MaFIA mice

Timed pregnancies were set up for MaFIA mice as described above. Fetal thymuses were harvested at E15.5 for FTOC. Briefly, culture wells were set up in a 12-well-plate 24 hour before FTOC. In each well, Whatman® Nuclepore™ Track-Etched membrane (WHA110409; Sigma) was placed on top of SURGIFORM® absorbable gelatin sponge (1974; Ethicon) in 1.5 mL of complete Dulbecco’s Modified Eagle Medium (DMEM) (supplemented with 10% Fetal Bovine Serum, 2 mM L-glutamine, 100 U/mL penicillin and 100 μg/mL streptomycin) (29,30). Three fetal thymuses were placed on top of the membrane for each well. B/B homodimerizer stock was diluted to 2.5 μM in complete DMEM and added to the thymus lobes at the time of culture setup and again on the following day. On day 2, the membrane with the thymus lobes was transferred to a fresh culture plate containing new gelatin sponge and complete DMEM. Cultures were maintained in 37 °C incubator until day 6, at which point the thymic lobes were processed for flow cytometry analysis as described above.

### Transcriptomic analysis

CD64^+^ cells were isolated from the thymus of C57BL/6J adult mice and processed for scRNA-seq as described below. Data from each scRNA-seq experiment underwent pre-processing as detailed below. Quality control filters were applied to remove cells with more than 12.5% mitochondrial gene expression, fewer than 500 reads, or more than 6,000 reads, ensuring the retention of high-quality cells for analysis. Macrophages and monocytes were identified using a module scoring approach, and their respective cell barcodes were recorded. These barcodes were used for subsetting before merging the datasets from the two scRNA-seq experiments with published TM data from Dzhagalov’s group. The merging was performed using Harmony to align the datasets and correct for batch effects. Following integration, cells were clustered, and module scores were assigned to all identified clusters. Non-relevant cell types, including B cells, T cells, and DCs, were excluded from subsequent analyses to focus on macrophages and monocytes.

### Analysis of scRNA-Seq and data integration

All analyses were performed using R software version 4.4.1. Cells expressing fewer than 500 genes or more than 6,000 genes were excluded from the dataset (GEO: GSE326594). Additionally, cells with greater than 12.5% mitochondrial gene content were removed to ensure data quality. The dimensionality of the dataset was assessed using an Elbow plot to determine the appropriate number of principal components. Gene ontology (GO) enrichment analysis of differentially expressed genes (DEGs) was conducted to identify overrepresented biological processes using the FindMarkers function from the Seurat package (31). The data were then integrated with previously published scRNA-seq datasets on TMs (GEO: GSE185460), as well as datasets from heart, liver, and lung macrophages (GEO: GSE188647), from the laboratories of Drs. Dzhagalov and Epelman, respectively. Batch correction across datasets was performed using the Harmony package (27,32,33). Following integration, the merged dataset was normalized using the SCTransform method to account for technical variation. For data visualization, Uniform Manifold Approximation and Projection (UMAP) was applied to reduce dimensionality and enable visualization of cellular clustering patterns.

### Trajectory analysis using Monocle3

Trajectory analysis was performed using Monocle3 to examine transcriptional relationships among monocytes and TMs (34). Following SCTransform normalization and integration in Seurat, the expression matrix, cell metadata, and gene metadata were used to construct a Monocle3 CellDataSet. Principal component analysis and UMAP were used for dimensionality reduction, followed by re-clustering and graph learning. *Ly6c2+ Ccr2+* monocytes were assigned as the root population based on their established precursor phenotype. Pseudotime therefore represents a model of transcriptional progression from a monocyte-like state and is interpreted together with, rather than independently of, the genetic fate-mapping and CCR2-dependence data.

### Statistical analysis

Data are presented as the mean ± standard error of the mean (SEM). Univariate comparisons between two groups were performed using the Mann-Whitney U test. For analyses involving more than two groups, the normality of the data was assessed using the D’Agostino-Pearson test and/or inspection of Q-Q plots, while homoscedasticity (equal variance) was evaluated using the Brown-Forsythe test or by examining residual plots. Non-parametric comparisons across multiple groups were conducted using the Kruskal-Wallis test, followed by post-hoc pairwise comparisons of the mean ranks of each group with the mean rank of the control group. Multiple comparisons were adjusted using Dunn’s test to control for type I error. For pairwise comparisons involving more than two groups, the Holm-Šídák method was applied to correct for multiple comparisons, reducing the likelihood of type I error. All statistical analyses were performed using GraphPad Prism version 10.

## RESULTS

### Identification of Two Distinct Tissue-Resident Macrophage Subsets in the Adult Thymus

To investigate the heterogeneity of phagocytic cells within the thymus, we performed multi-parameter flow cytometry on hematopoietic cells (CD45^+^) from the thymus of adult mice. We first identified the total TM population based on the co-expression of the canonical markers CD64 and F4/80 (**Figure 1A-B**). The majority of these CD64^+^ F4/80^+^ cells also expressed the phagocytic receptor MerTK, consistent with their identity as macrophages (35).

**Figure 1.**
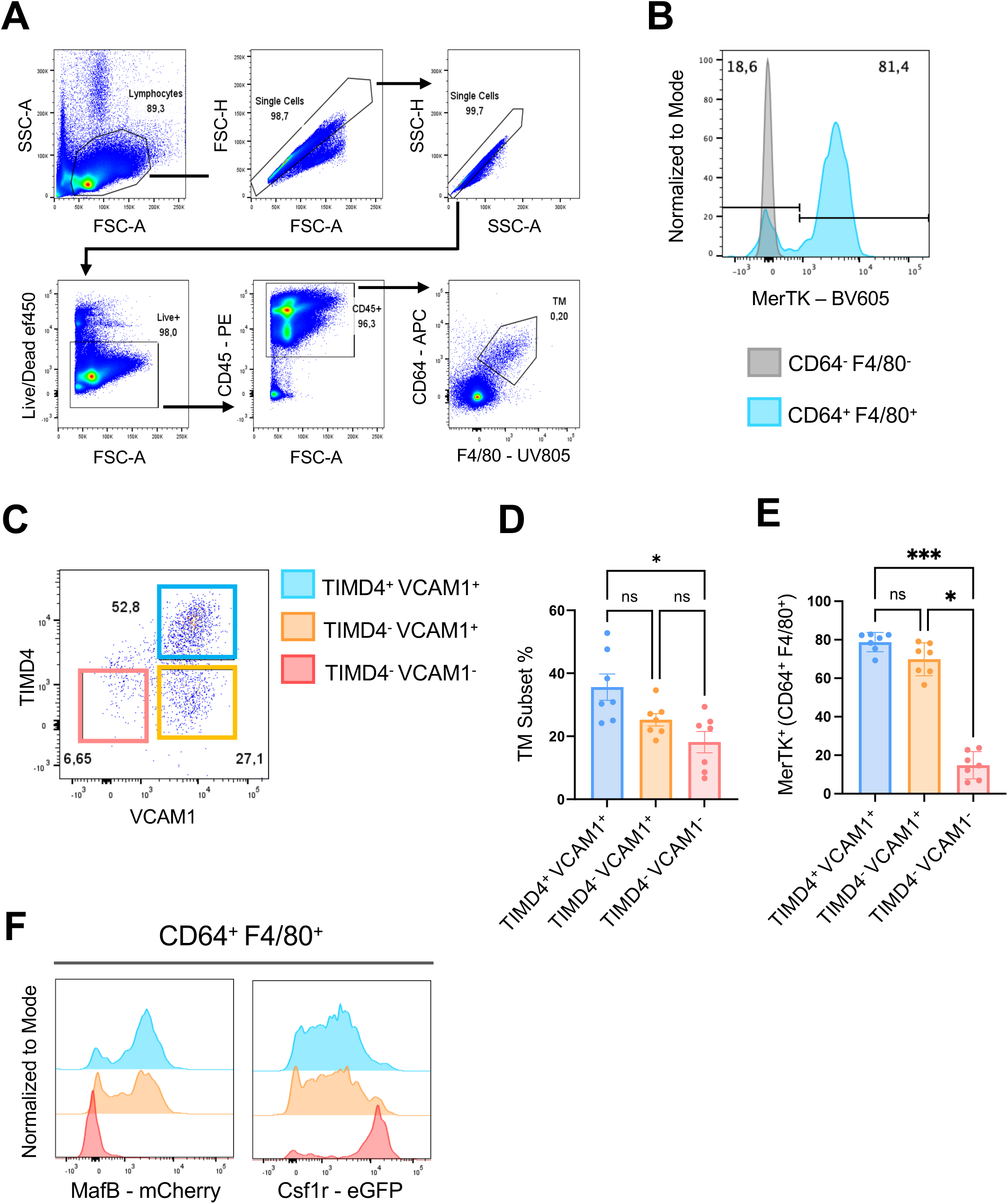
Flow-cytometric characterization of thymic macrophage populations. Thymuses from adult C57BL/6 mice were enzymatically digested and analyzed by flow cytometry. A) Gating strategy for CD64+ F4/80+ TMs. B) Representative MerTK histogram for TMs (blue) and CD64-F4/80-lymphocytes (red). C) TIMD4/VCAM1 definition of TIMD4+ VCAM1+, TIMD4-VCAM1+, and TIMD4-VCAM1-populations. D) Frequencies of the three TM populations. E) MerTK+ frequencies. F) MafB-mCherry and *Csf1r*-EGFP reporter expression evaluated within the same TIMD4/VCAM1 gates defined in panel C. Data are mean ± SEM (n = 7 per group from two independent experiments). Kruskal-Wallis test (*p ≤ 0.05).

Analysis of TIMD4 and VCAM1 expression resolved three populations within the CD64+ F4/80+ gate: a major TIMD4+ VCAM1+ population, a smaller TIMD4-VCAM1+ population, and a minor TIMD4-VCAM1-population (**Figure 1C-E**). The two VCAM1+ gates correspond phenotypically to the TIMD4+ cortical and TIMD4-/CX3CR1+ medullary/cortico-medullary macrophage populations described by Zhou *et al*., while VCAM1 provides an additional prospective discriminator (27). Both VCAM1+ populations expressed the macrophage-associated MafB-mCherry reporter. By contrast, the TIMD4-VCAM1-population was MafB-low/negative and displayed high *Csf1r*-EGFP, consistent with a monocyte-enriched phenotype (**Figure 1F**). These results establish a flow-cytometric framework that links our prospectively defined populations to the major TM states identified transcriptionally and anatomically in prior work (27).

Short-term intravenous (i.v.) labeling with anti-CD45 demonstrated that most TIMD4+ VCAM1+ and TIMD4-VCAM1+ cells were shielded from the circulation, supporting their predominantly parenchymal localization (**Figure 2A-C**). The broad Ly6C+ CD11b+ comparison gate contained both iv-labeled and unlabeled cells, whereas the more stringently defined TIMD4-VCAM1-gate within CD64+ F4/80+ cells contained a smaller i.v.-labeled fraction. The two VCAM1+ populations were Ly6C-low/negative and expressed substantially less CCR2 than TIMD4-VCAM1-cells. CX3CR1 expression was enriched in the TIMD4-VCAM1+ population relative to TIMD4+ VCAM1+ macrophages (**Figure 2D-E**), further aligning these populations with the TIMD4+ and CX3CR1+ TM states described previously (27). In contrast, the TIMD4-VCAM1-population was enriched for Ly6C, CCR2, and CX3CR1, supporting its designation as a monocyte-enriched thymic population.

**Figure 2.**
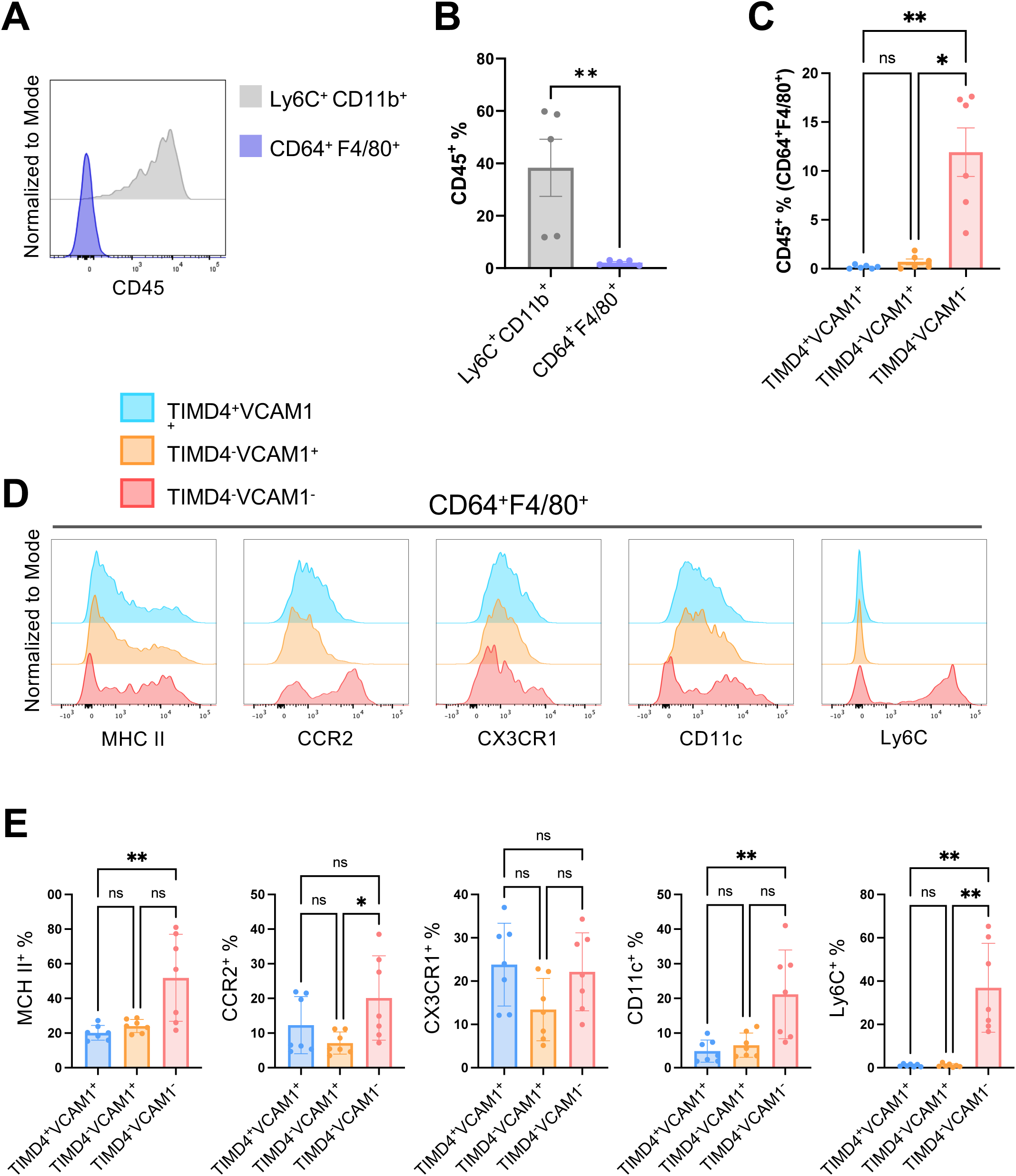
Intravascular labeling and monocyte-associated marker expression distinguish thymic myeloid populations. A) Short-term intravenous (i.v.) anti-CD45 labeling identifies blood-exposed cells. B) Frequencies of i.v. CD45-labeled cells in the broad Ly6C+ CD11b+ gate and the CD64+ F4/80+ macrophage gate. The Ly6C+ CD11b+ gate contains heterogeneous blood-exposed and parenchymal cells and is not equivalent to the TIMD4-VCAM1-gate. C) i.v. CD45 labeling within TIMD4/VCAM1-defined populations. D-E) Relative Ly6C, CCR2, and CX3CR1 signal across the three populations; descriptions are comparative rather than categorical. Data are mean ± SEM (n = 5-7 per group from two independent experiments). Mann-Whitney test for two-group comparisons and Kruskal-Wallis test for three-group comparisons (*p ≤ 0.05; **p ≤ 0.01).

Together, these data identify two predominantly parenchymal VCAM1+ TM populations and distinguish them from a TIMD4-VCAM1-monocyte-enriched population. This gating framework connects surface phenotype, anatomical accessibility, and reporter expression, enabling prospective analysis of the major TM populations.

### Transcriptomic Profiling by scRNA-seq Confirms Two Functionally Distinct Macrophage Subsets

To gain deeper insight into the transcriptional identities of these macrophage populations, we performed scRNA-seq on sorted thymic CD45+ CD64+ cells. Unbiased clustering of the resulting 3,153 cells revealed five distinct myeloid clusters, including monocytes, proliferating macrophages, and two major quiescent macrophage populations that corresponded to our flow-cytometric gates (**Figure 3A**).

**Figure 3.**
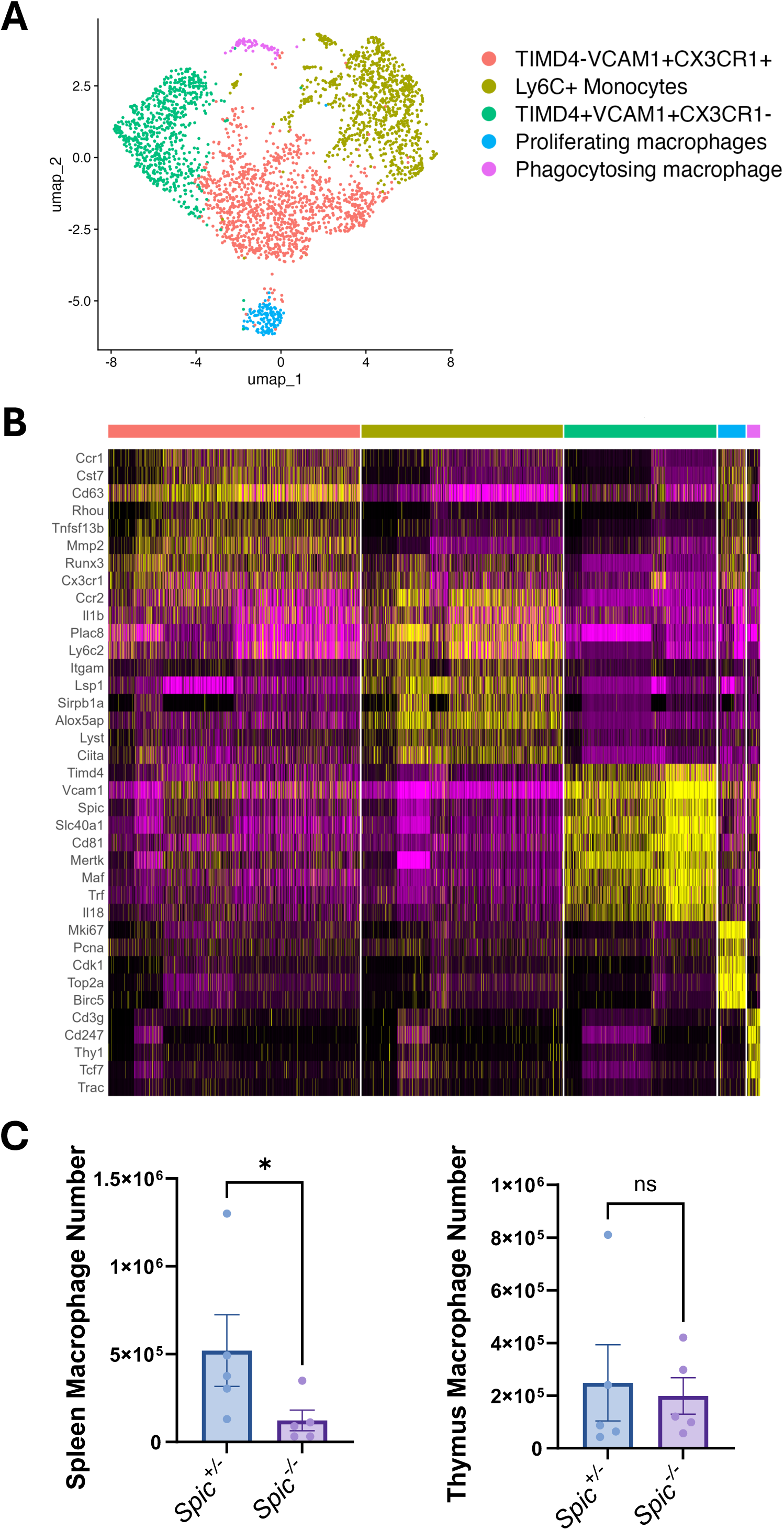
Single-cell transcriptomic profiles of thymic macrophages. CD64+ cells from six-week-old C57BL/6 thymuses were analyzed by 10X Genomics 3’ scRNA-seq in two independent experiments, each pooling six mice, and integrated with the Zhou *et al*. dataset using Harmony. A) UMAP showing five annotated myeloid clusters; panel colors denote cluster identity. B) Heatmap of row-scaled differentially expressed genes; yellow and purple represent higher and lower relative scaled expression, respectively, and are not equivalent to flow-cytometric fluorescence intensity. C) Total CD64+ F4/80+ macrophage numbers in thymus and spleen of *Spic-/-* and control mice; TM subpopulations were not separately quantified in this panel. Data are mean ± SEM (n = 4-5 per group from two independent experiments), Mann-Whitney test (*p ≤ 0.05).

One major macrophage cluster expressed *Timd4*, *Vcam1*, *Mertk*, and *Maf*, corresponding to the TIMD4+ VCAM1+ population (**Figure 3B**). A second cluster expressed *Vcam1* and *Cx3cr1*, aligning with the TIMD4-VCAM1+ population (**Table 1**). The TIMD4+ VCAM1+ cluster also expressed *Spic*, the lineage-defining transcription factor required for splenic red pulp macrophages (RPMs) (36). We therefore examined the TM compartment in *Spic-/-* mice. As expected, splenic RPMs were markedly reduced, whereas total CD64+ F4/80+ TM abundance was preserved (**Figure 3C**). Thus, despite sharing *Spic* expression with RPMs, maintenance of the overall TM compartment follows a distinct developmental program that does not require SpiC.

**Table 1.** Differentially expressed genes (DEGs) from scRNA-seq analysis. List of DEGs with an average log fold change > 1 and an adjusted p-value < 0.01, ordered by adjusted log fold change.

| **Ly6C<sup>+</sup> Monocytes** |  | **TIMD4<sup>+</sup> VCAM1<sup>+</sup> CX3CR1<sup>-</sup>** |  | **TIMD4<sup>-</sup> VCAM1<sup>+</sup> CX3CR1<sup>+</sup>** |  |
| --- | --- | --- | --- | --- | --- |
| *Lsp1* | *Ahnak* | *Cd163* | *Ccl2* | *Gpnmb* | *Lhfpl2* |
| *Sirpb1a* | *Cytip* | *Mrc1* | *Ccl4* | *Tnfsf13b* | *Chst2* |
| *Gm34084* | *Ifitm2* | *Mertk* | *Adgre1* | *Lhfpl2* | *Mmp2* |
| *Alcam* | *S100a11* | *Spic* | *Adgre2* | *Chst2* | *Pmepa1* |
| *Cbfa2t3* | *Ramp1* | *Timd4* | *Hmox1* | *Mmp2* | *Oas3* |
| *Sirpb1b* | *Lyst* | *Vcam1* | *Tlr7* | *Pmepa1* | *Cst7* |
| *Cnn2* | *Ccl6* | *Clec1b* | *Axl* | *Oasl1* | *Zmynd15* |
| *Samsn1* | *Itga4* | *Stab2* | *Fgr* | *Cst7* | *H2-Q7* |
| *Itgam* | *Il17ra* | *Fcgrt* | *Ccl8* | *Zmynd15* | *Ccr5* |
| *Alox5ap* | *Ccr2* | *Tgm2* | *Id2* | *H2-M2* | *H2-T22* |
| *Emb* | *Ly6c2* | *P2ry12* | *Sirpa* | *Sh3pxd2b* | *Bst2* |
| *Traf1* | *Il1b* | *Cd86* | *Runx1* | *Kctd14* | *Stat2* |
| *Dpp4* | *Batf3* | *Tlr4* | *Stat5b* | *Ifit1* | *Cebpa* |
| *S100a10* | *Rbpj* | *Il18* | *Cd84* | *Fam20c* | *Il1rn* |
| *Plac8* |  | *Ptgs1* | *Mapk14* | *Cp* | *Cxcl16* |
| *Stap1* |  | *Pparg* |  | *Spsb1* |  |
| *Sirpb1c* |  | *Adamdec1* |  | *Gpnmb* |  |
| *Rnase6* |  | *Cd38* |  | *Tnfsf13b* |  |

Gene Ontology analysis revealed pronounced functional specialization between the two TM populations. TIMD4-VCAM1+ CX3CR1+ TMs were enriched for antigen processing and presentation via MHC class I and for interferon-response pathways, consistent with a potential role in immune surveillance and T cell education (**Figure 4A**). Conversely, TIMD4+ VCAM1+ TMs were strongly enriched for phagocytosis and endocytosis pathways, supporting specialization for efferocytic clearance of apoptotic thymocytes (**Figure 4B**). Individual genes contributing to these subset signatures, including H2-K1, H2-D1, and Runx3, are reported in **Table 1**.

**Figure 4.**
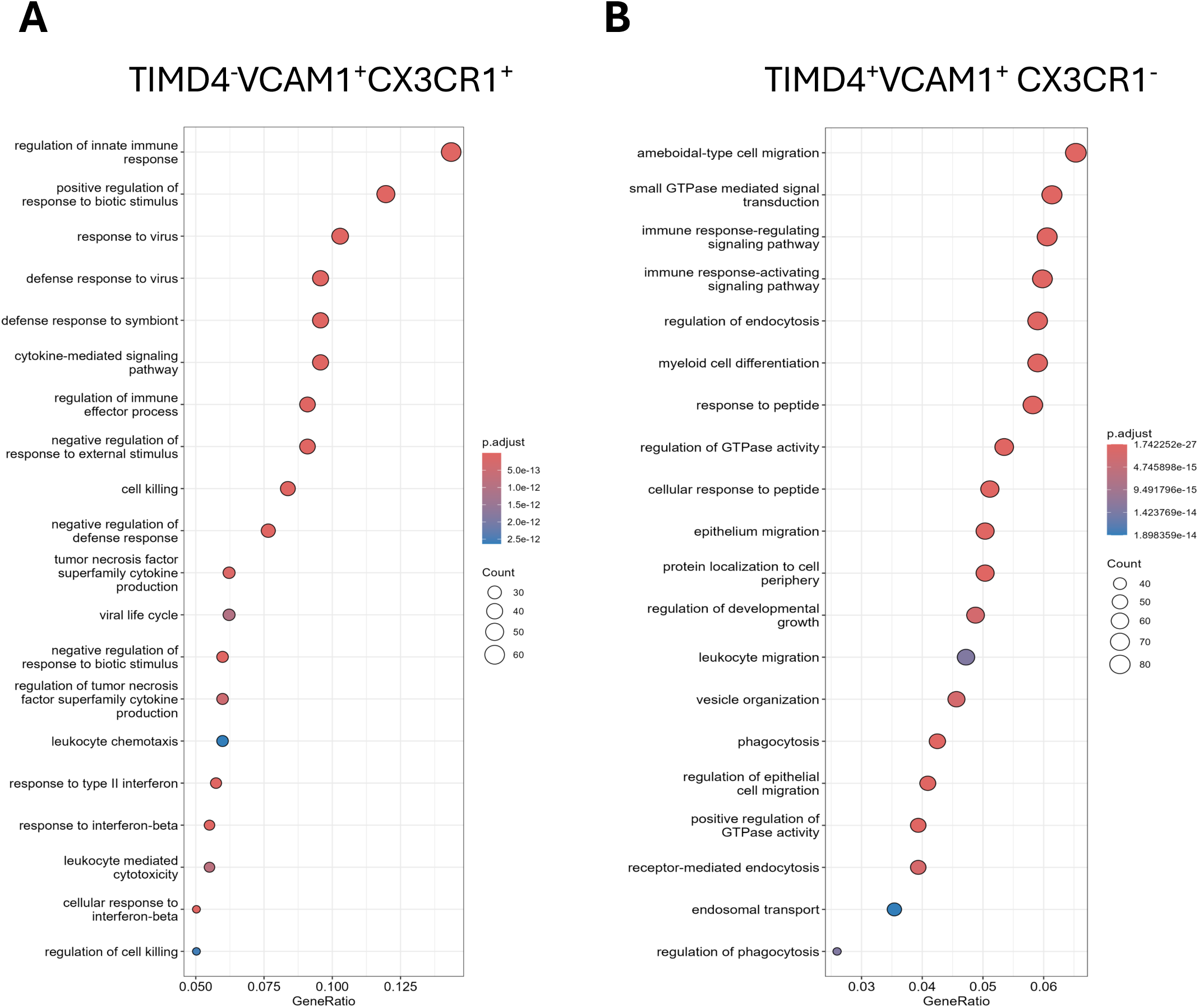
Gene Ontology pathway analysis of TM populations. A) Top enriched pathways in TIMD4-VCAM1+ CX3CR1+ TMs. B) Top enriched pathways in TIMD4+ VCAM1+ TMs. The panels display pathway-level enrichment; individual genes mentioned in the text are reported in Table 1 and are not plotted separately in this Figure.

A subcluster within the TIMD4+ VCAM1+ population co-expressed macrophage genes together with thymocyte-associated transcripts, including *Cd3g* and *Trac* (**Figure 3B**). In the context of the strong phagocytosis and endocytosis program in this population, the presence of thymocyte-derived transcripts is consistent with ongoing uptake of thymocyte material and provides additional support for efferocytic specialization. As with other phagocyte single-cell datasets, contributions from ambient RNA or rare multiplets cannot be fully distinguished from engulfed RNA, and we therefore interpret this signature together with the broader pathway evidence.

Exploratory Monocle3 analysis, rooted in *Ly6c2*+ *Ccr2*+ monocytes, placed monocytes and macrophage states along a connected transcriptional manifold. The ordering was consistent with progressive acquisition of macrophage-associated programs from a monocyte-like state and provided a transcriptional complement to the fate-mapping and CCR2-dependence experiments. As pseudotime models transcriptional relationships rather than prospective cell fate, it only allows us to predict a potential conversion between the two mature TM populations.

### The Thymic Macrophage Compartment Undergoes Dynamic Remodeling During Development

To understand how the TM compartment evolves in parallel with the thymus itself, we analyzed the number and composition of TM subsets during ontogeny, from embryonic development through adulthood. Consistent with known thymic biology, the absolute number of total thymocytes and TMs peaked in young, 3-week-old mice, before declining with age, indicating that the size of the TM pool is coupled to the overall cellularity of the organ (**Supplemental Figure 1A-B**).

TM composition was extensively remodeled across development. At embryonic day 15 (E15), the thymic myeloid compartment was dominated by TIMD4-VCAM1-cells expressing CD11b and CCR2, consistent with recently recruited or immature myeloid precursors (**Figure 5A**).

**Figure 5.**
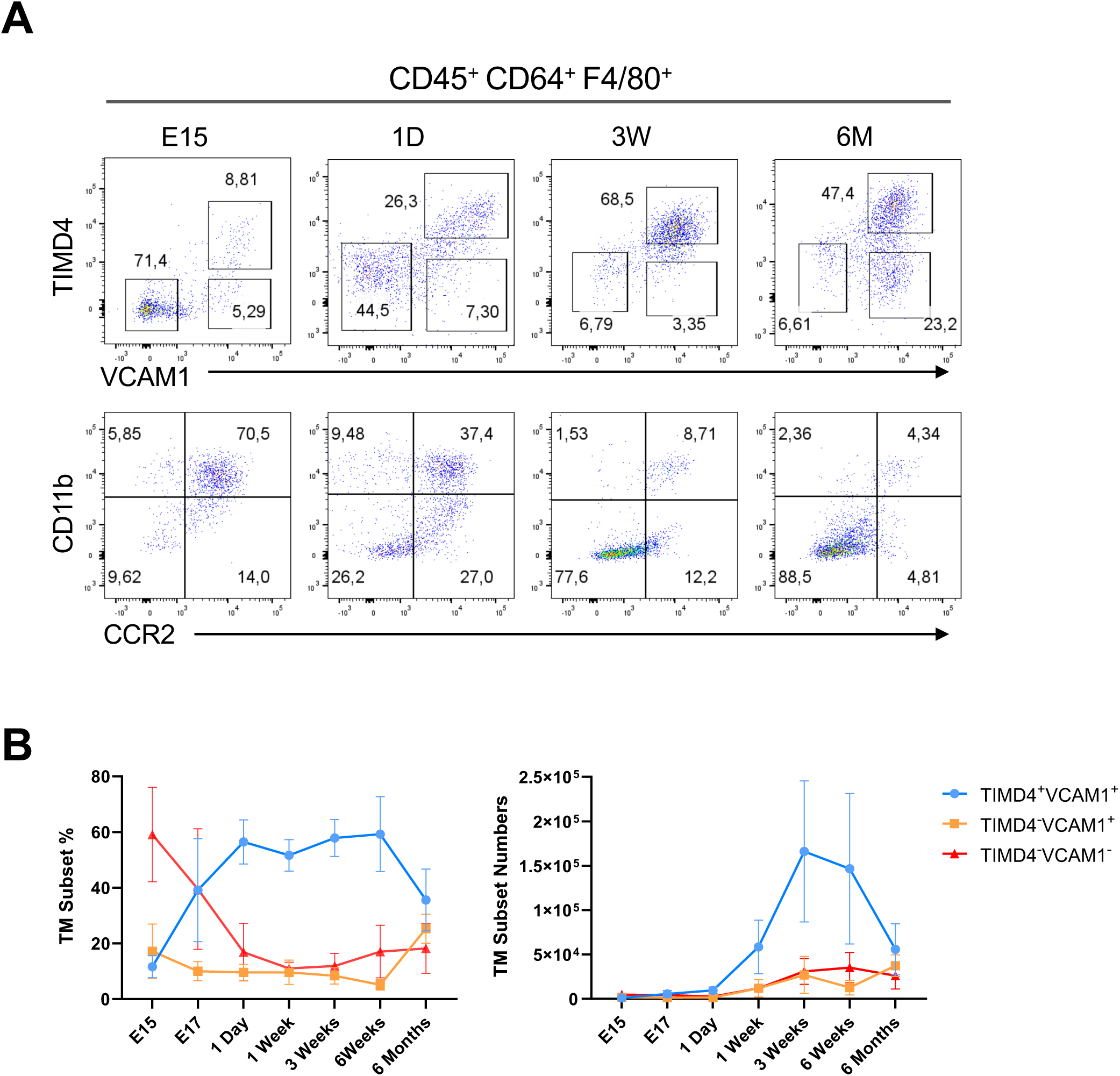
Developmental progression of TM subsets. Thymuses were harvested from mice at various developmental time points and enzymatically digested to prepare single-cell suspensions. Cells were stained with antibodies for flow cytometry analysis. A) Expression of TM markers across different age groups. B) Comparison of the numbers and percentages of TM in mice at different ages. Data are presented as mean ± SEM (n = 4-12 per group from 2-4 independent experiments).

Following birth, TIMD4+ VCAM1+ macrophages expanded rapidly and became the dominant TM population during the period of maximal thymic growth (**Figure 5B**). With the onset of age-associated thymic involution, the proportion of TIMD4+ VCAM1+ macrophages declined, whereas TIMD4-VCAM1+ macrophages progressively increased. These data reproduce the age-associated shift reported by Zhou *et al*. and show that our VCAM1-based gating strategy captures the developmental remodeling of the two major TM populations (27).

Thus, the thymic macrophage compartment is developmentally dynamic: immature myeloid cells predominate in the fetal organ, TIMD4+ VCAM1+ macrophages expand with postnatal thymic growth, and TIMD4-VCAM1+ macrophages become increasingly prominent during involution. The changing abundance of these populations establishes the developmental context for their distinct ontogenetic contributions.

### Thymic Macrophage Subsets Arise from Distinct Developmental Origins

Flt3-Cre × Rosa-mTmG fate mapping revealed unequal developmental contributions to the two TM populations. In this model, cells with a history of FLT3 expression switch from membrane Tomato to membrane GFP, whereas FLT3-independent cells remain Tomato+ (**Figure 6A**) (28). TIMD4+ VCAM1+ macrophages contained a large Tomato+ fraction, demonstrating a substantial contribution from FLT3-independent embryonic/fetal precursors. In contrast, TIMD4-VCAM1+ macrophages and thymic monocytes were predominantly GFP+, consistent with derivation from FLT3-expressing hematopoietic progenitors (**Figure 6B-C**). Tomato+ GFP+ events likely represent ongoing reporter transition or persistence of stable membrane Tomato after recombination and were included within the overall assessment of reporter history rather than treated as a separate lineage.

**Figure 6.**
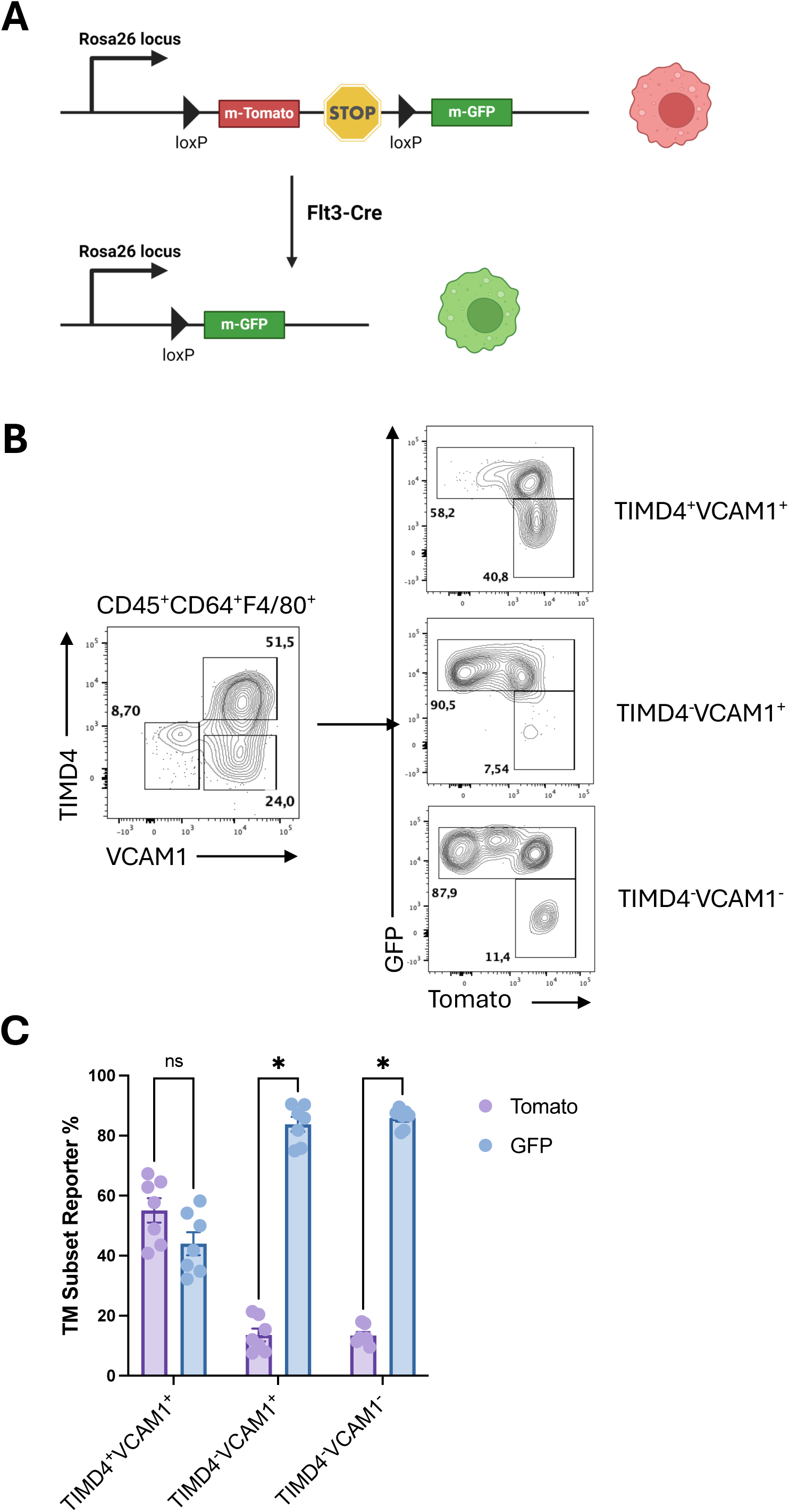
Flt3-Cre reporter history in thymic myeloid populations. A) In Flt3-Cre × Rosa-mTmG mice, Cre-mediated recombination switches membrane Tomato expression to membrane GFP in cells with *Flt3* expression history. B) Representative Tomato/GFP profiles within TIMD4+ VCAM1+, TIMD4-VCAM1+, and TIMD4-VCAM1-gates. Tomato+ GFP+ events are interpreted as reporter-transition/persistence events and not as a distinct lineage. C) Reporter frequencies among the three populations. CX3CR1 phenotype is defined independently in Figure 2 and is not displayed in this Figure. Data are mean ± SEM (n = 7 per group from two independent experiments). Mann-Whitney tests with Holm-Šídák correction (***p ≤ 0.001).

Because TIMD4-VCAM1+ macrophages showed greater FLT3-lineage history than their TIMD4+ VCAM1+ counterparts, we asked whether maintenance of this population depends on CCR2-mediated precursor recruitment. We therefore analyzed thymic myeloid populations in *Ccr2* knockout (*Ccr2-/-*) mice. Overall thymocyte cellularity and the abundance of TIMD4+ VCAM1+ macrophages were preserved, whereas both TIMD4-VCAM1+ macrophages and the TIMD4-VCAM1-monocyte-enriched population were significantly reduced (**Figure 7A-D**). This selective effect links CCR2-dependent recruitment to maintenance of the TIMD4-VCAM1+ arm of the thymic macrophage network.

**Figure 7.**
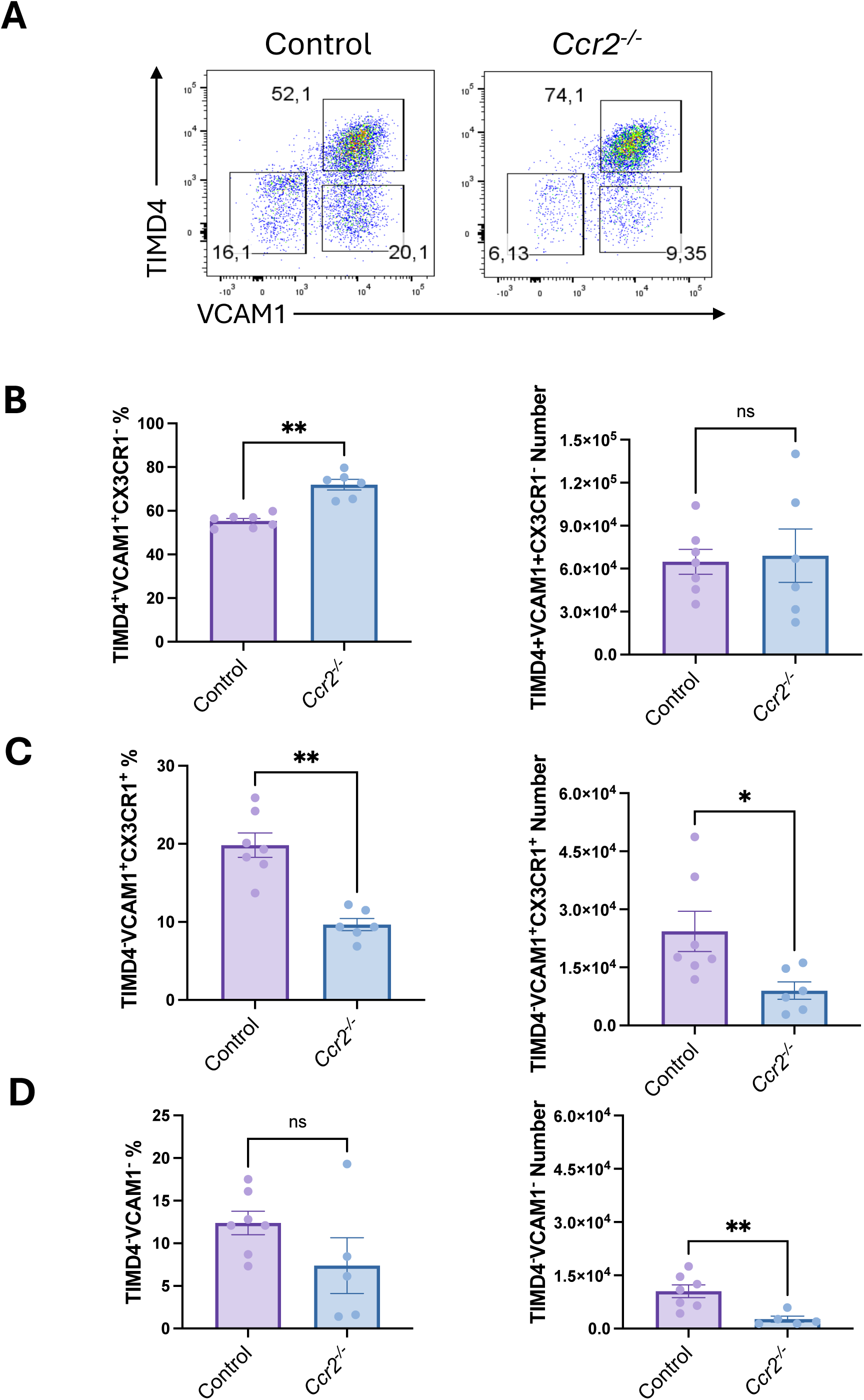
CCR2 dependence of TIMD4/VCAM1-defined thymic myeloid populations. A) Representative TIMD4/VCAM1 profiles from *Ccr2-/-* and control mice. B-D) Frequencies and absolute numbers of TIMD4+ VCAM1+, TIMD4-VCAM1+, and TIMD4-VCAM1-populations. CX3CR1 phenotype is defined independently in Figure 2 and is not displayed in this Figure. Data are mean ± SEM (n = 6-7 per group from two independent experiments). Two-tailed Mann-Whitney test (*p ≤ 0.05; **p ≤ 0.01; ***p ≤ 0.001).

Together, the fate-mapping and *Ccr2-/-* data support a dual-contribution model for the TM network. TIMD4+ VCAM1+ macrophages retain a prominent FLT3-independent embryonic/fetal contribution, whereas TIMD4-VCAM1+ macrophages show greater FLT3 history and depend on CCR2-mediated recruitment. The selective reduction of TIMD4-VCAM1+ macrophages and thymic monocytes in *Ccr2-/-* mice provides functional evidence that circulating precursors contribute to maintenance of this arm of the TM compartment.

### *Csf1r*-Expressing Myeloid Cells Support Thymocyte Progression Across the DN3-to-DN4 Checkpoint

To determine whether *Csf1r*-expressing myeloid cells contribute to early T cell development, we used the MaFIA (*Csf1r*-EGFP-NGFR/FKBP1A/TNFRSF6) model, in which AP20187 dimerizes an FKBP-Fas suicide protein and induces apoptosis in transgene-expressing cells (**Supplemental Figure 2A-B**) (37). Both TM populations and thymic monocytes expressed the *Csf1r*-EGFP reporter (**Supplemental Figure 2C-E**), making this system well suited to perturb the thymic myeloid compartment. We applied this experimental approach to E15.5 fetal thymic organ culture (FTOC), which preserves the three-dimensional thymic microenvironment while avoiding systemic consequences of myeloid-cell depletion (38), and treated FTOCs from MaFIA mice with AP20187 at culture initiation (**Figure 8A**).

**Figure 8.**
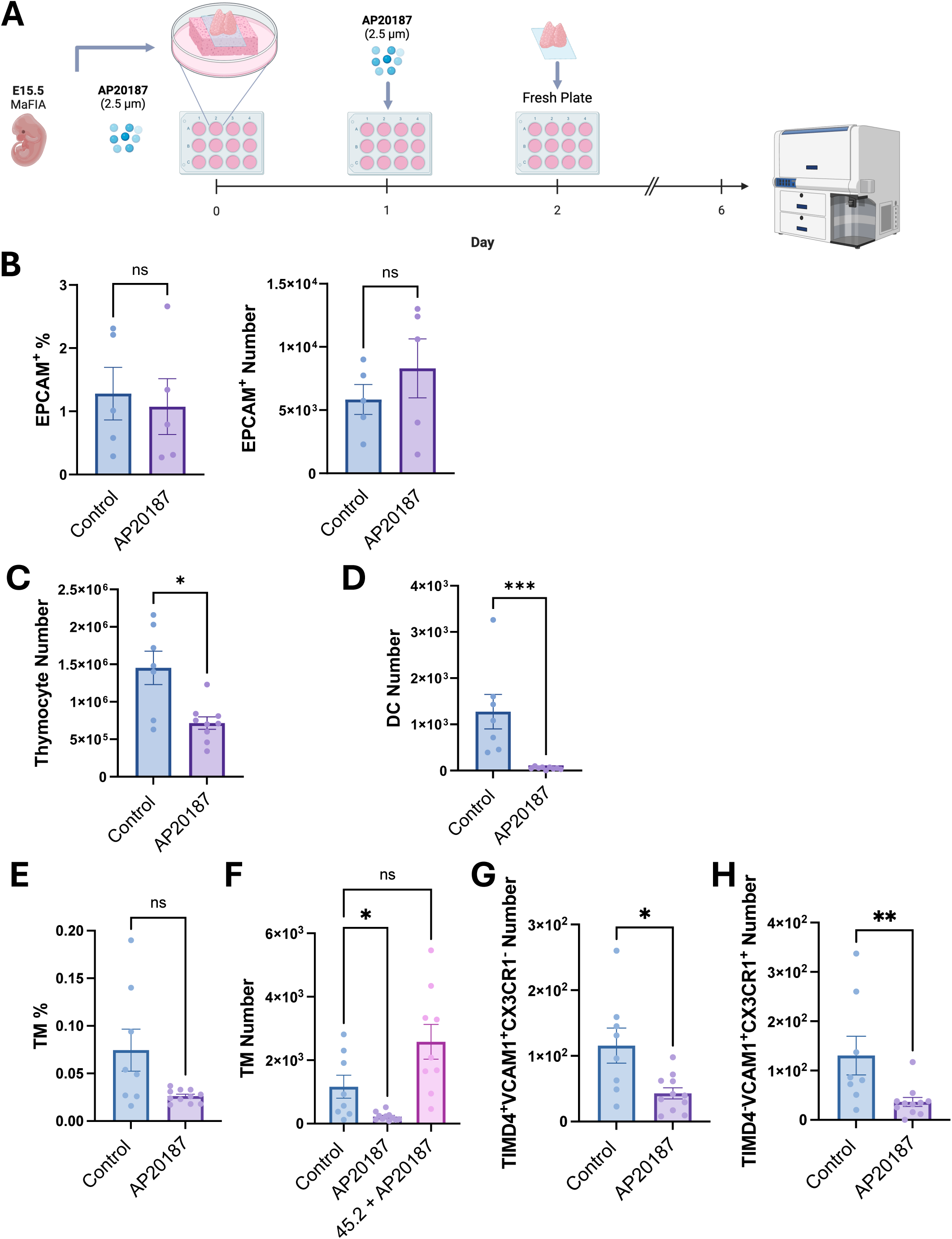
Efficient depletion of *Csf1r*-expressing myeloid cells in MaFIA fetal thymic organ culture. E15.5 MaFIA thymus lobes were treated with AP20187 at culture initiation and the following day, transferred to fresh FTOC media on day 2, and analyzed on day 6. A) Experimental workflow. B) EpCAM+ CD45-thymic epithelial-cell frequencies and numbers. C) Total thymocyte numbers. D) CD45+ CD11c+ MHC-II^high^ DC numbers. E-F) TM frequencies and absolute numbers in untreated and AP20187-treated MaFIA FTOCs and in AP20187-treated C57BL/6J (CD45.2) control FTOCs. CD45.2 denotes the congenic allele of the C57BL/6J control. G-H) Numbers of TIMD4+ VCAM1+ and TIMD4-VCAM1+ TMs. The preservation of epithelial-cell and control-FTOC TM numbers, together with depletion of both MaFIA TM populations, demonstrates transgene-dependent perturbation of the *Csf1r*-expressing thymic myeloid compartment. Data are mean +/− SEM (n = 8-11 per group from three independent experiments). Mann-Whitney or Kruskal-Wallis tests as indicated (*p <= 0.05; **p <= 0.01; ***p <= 0.001).

AP20187 treatment preserved the EpCAM+ CD45-thymic epithelial-cell compartment (**Figure 8B**) while markedly reducing total TMs and both TIMD4/VCAM1-defined TM populations (**Figure 8E-H**). Importantly, TM abundance was unchanged in AP20187-treated C57BL/6 control FTOCs lacking the MaFIA transgene (**Figure 8F**), demonstrating that depletion required the transgenic suicide system rather than exposure to the dimerizer alone. Total thymocyte and CD11c+ MHC-II^high^ DC numbers were also reduced (**Figure 8C-D**). These results establish efficient, transgene-dependent perturbation of the *Csf1r*-expressing thymic myeloid compartment within an epithelial environment that remained numerically intact.

Depletion of *Csf1r*-expressing cells impaired αβ T-lineage cell development, leading to a marked reduction in CD4+ CD8+ double-positive (DP) thymocytes, whereas γδ T cell output was preserved (**Figure 9A-D**). To localize the αβ lineage developmental defect, we examined upstream DN populations (**Figure 10A**). AP20187-treated MaFIA FTOCs showed a coordinated accumulation of DN3 (CD44-CD25+) cells and reduction of DN4 (CD44-CD25-) cells (**Figure 10B**). The reciprocal change in sequential developmental populations identifies the DN3-to-DN4 transition as the principal stage affected by the loss of *Csf1r*-expressing cells.

**Figure 9.**
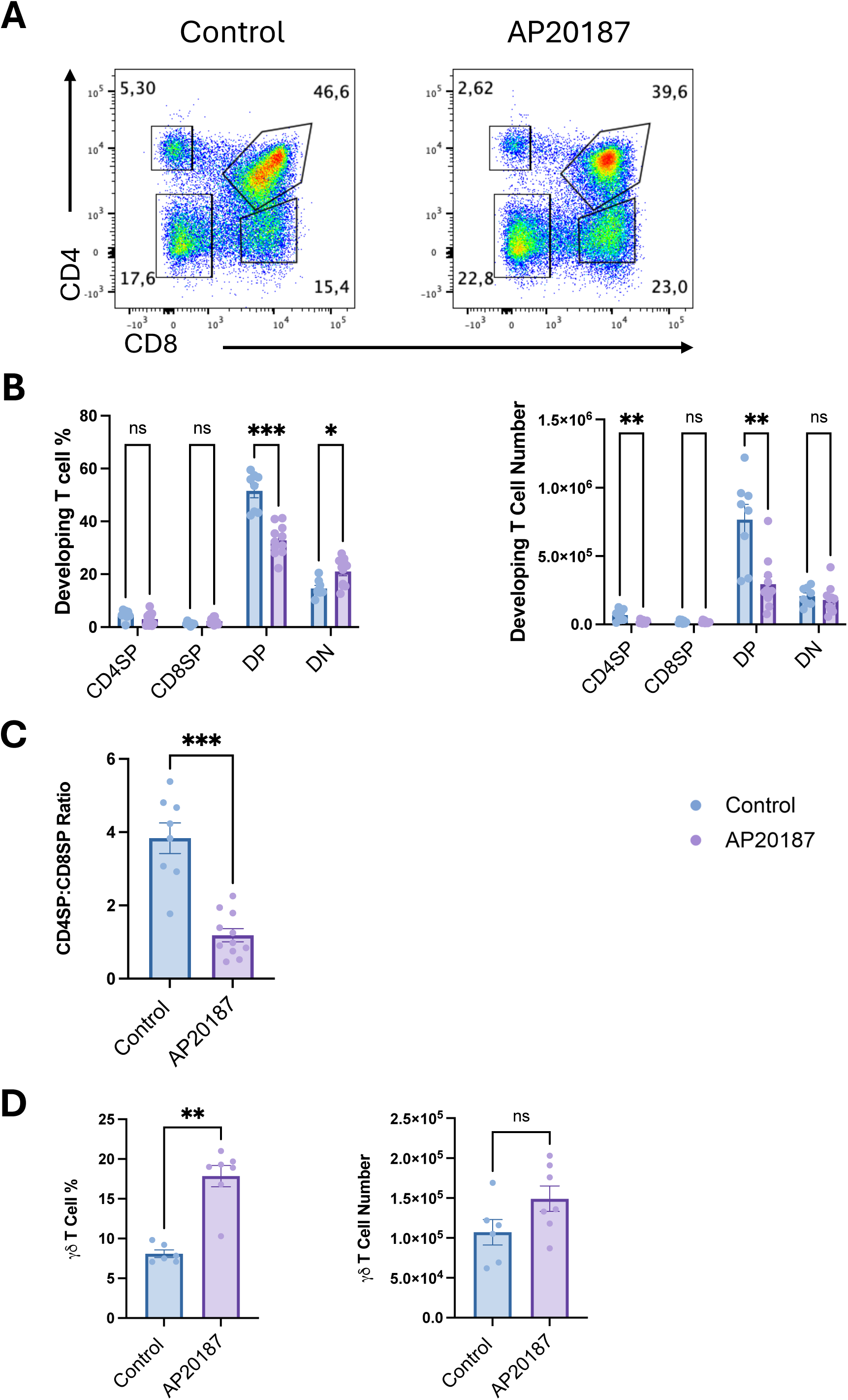
*Csf1r*-expressing myeloid cells support alpha-beta T cell development in FTOC. A) Representative CD4/CD8 profiles. B) Frequencies and absolute numbers of thymocyte populations in untreated and AP20187-treated MaFIA FTOCs. C) CD3+ CD4SP-to-CD3+ CD8SP ratio. D) Frequencies and numbers of γδ T cells. Depletion reduced αβ-lineage DP thymocyte production while preserving γδ T cell output. Data are mean +/− SEM (n = 8-11 per group from three independent experiments; γδ T cells, n = 6-7 from two experiments). Mann-Whitney tests with Holm-Sidak correction (*p <= 0.05; **p <= 0.01; ***p <= 0.001).

**Figure 10.**
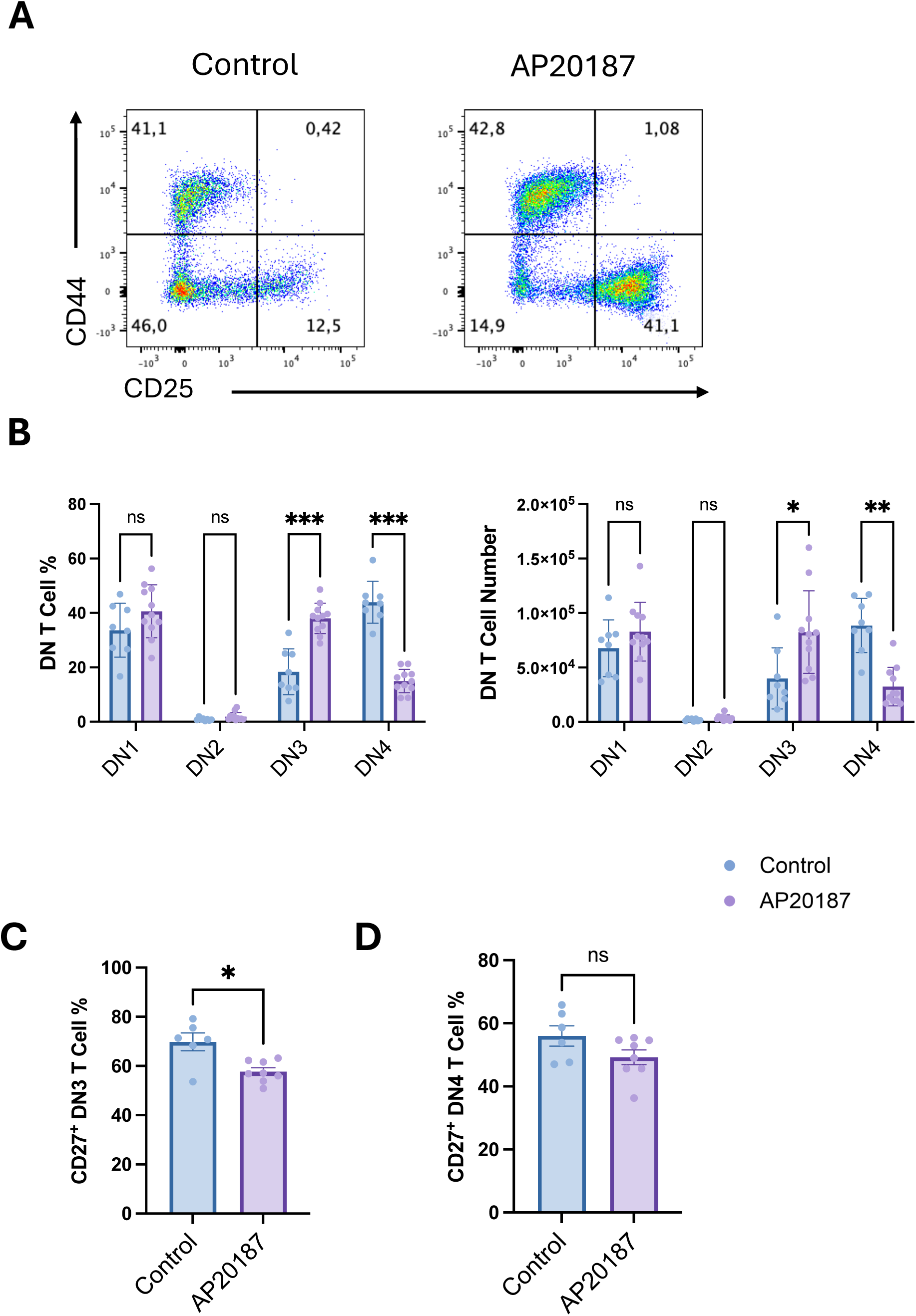
*Csf1r*-expressing myeloid cells support progression across the DN3-to-DN4 checkpoint. A) Representative CD44/CD25 profiles of DN populations. B) Frequencies and absolute numbers of DN subsets in untreated and AP20187-treated MaFIA FTOCs. C-D) Quantified frequencies of CD27+ DN3 and DN4 cells. The coordinated accumulation of DN3 cells, reduction of DN4 cells, and lower CD27 expression identify impaired progression at the β-selection transition. Data are mean +/− SEM (DN subsets, n = 8-11 from three independent experiments; CD27, n = 6-8 from two experiments). Mann-Whitney tests with Holm-Sidak correction (*p <= 0.05; **p <= 0.01; ***p <= 0.001).

We next examined CD27 expression, which is upregulated as DN3 thymocytes successfully receive pre-TCR-associated signals and progress through β-selection. The frequency of CD27+ DN3 cells was significantly reduced in AP20187-treated MaFIA FTOCs (**Figure 10C-D**). Considered together, DN3 accumulation, DN4 loss, reduced CD27 expression, and diminished DP cell output, we have strong evidence that *Csf1r*-expressing myeloid cells support thymocyte differentiation at the β-selection checkpoint. Thus, the thymic myeloid compartment contributes not only to tissue clearance and homeostasis but also to the microenvironmental conditions required for efficient early αβ T cell development.

## DISCUSSION

This study defines the mouse TM compartment as a developmentally dynamic network of phenotypically and transcriptionally specialized macrophages and connects the thymic myeloid niche to an early checkpoint in αβ T cell development. By integrating prospective surface-marker definition with single-cell transcriptomics, fate mapping, genetic perturbation, and organ culture, we identify two major TM populations with distinct functional programs and developmental contributions, demonstrate CCR2 dependence of the TIMD4-VCAM1+ population, and show that *Csf1r*-expressing myeloid cells support efficient DN3-to-DN4 progression.

The TIMD4+ VCAM1+ population displayed a coherent efferocytic identity, including Mertk expression, enrichment of phagocytosis and endocytosis pathways, and detection of thymocyte-associated transcripts within a macrophage subcluster. Although these cells expressed *Spic*, preservation of total TMs in *Spic-/-* mice distinguishes maintenance of the thymic compartment from the SpiC-dependent program of splenic red pulp macrophages (36). In contrast, TIMD4-VCAM1+ macrophages were enriched for antigen-presentation and interferon-response pathways, suggesting specialization for immune surveillance and interactions with developing thymocytes. The surface-marker correspondence between these transcriptional states and the cortical TIMD4+ and medullary/cortico-medullary CX3CR1+ populations described by Zhou *et al.* provides a unified phenotypic and functional framework (27).

The ontogeny experiments further reveal how this functional diversity is maintained. TIMD4+ VCAM1+ macrophages retained a substantial FLT3-independent embryonic/fetal contribution, whereas TIMD4-VCAM1+ macrophages showed greater FLT3 history. The selective loss of TIMD4-VCAM1+ macrophages and thymic monocytes in *Ccr2-/-* mice provides functional support for continuing recruitment from circulating precursors. These results confirm the broad dual-origin framework established by Zhou *et al*. (27), and extend it by identifying CCR2 as a requirement for maintenance of the TIMD4-VCAM1+ arm of the network under steady-state conditions.

The functional studies reveal that the early αβ T cell program is sensitive to disruption of the thymic myeloid niche. The combination of DN3 accumulation, DN4 reduction, diminished CD27 expression, and reduced DP output localizes this requirement to progression at or near β-selection (41,42). Preservation of γδ T cells and thymic epithelial cell numbers argues against a uniform collapse of fetal thymic development and instead identifies a selective vulnerability of the αβ-lineage developmental pathway. These findings expand the established homeostatic role of thymic myeloid cells by showing that their presence is linked to efficient navigation of the first TCRβ rearrangement-dependent checkpoint.

Several complementary mechanisms could account for this myeloid-cell requirement. TIMD4+ macrophages are positioned and transcriptionally equipped to clear apoptotic thymocytes efficiently; their depletion could increase local cellular debris or alter the anti-inflammatory signals normally generated during efferocytosis, including TGF-β and IL-10 (51–53). Myeloid cells may also provide survival, metabolic, or receptor-ligand signals that cooperate with pre-TCR and Notch pathways necessary for DN3-to-DN4 progression (43–48). The MaFIA mouse model depletes a *Csf1r*-expressing compartment that includes TMs, monocytes, and DCs, and our experiments did not directly assess *Csf1r*-transgene expression or AP20187-induced apoptosis in purified DN thymocytes. Consequently, the current data establish the requirement at the level of the *Csf1r*-expressing myeloid niche rather than assigning it to one macrophage subset or a single molecular mechanism. This evidence-matched cellular resolution sharpens the developmental conclusion and provides a focused basis for future cell-restricted studies.

In conclusion, this study provides an integrated phenotypic, transcriptional, and developmental map of mouse TMs and demonstrates that the TIMD4-VCAM1+ population depends on CCR2-mediated precursor input. Most importantly, the organ culture experiments identify an intact *Csf1r*-expressing myeloid compartment as a requirement for efficient progression across the DN3-to-DN4 checkpoint. These findings place thymic macrophage heterogeneity within a broader developmental framework and establish the thymic myeloid niche as an active component of early T cell development.

## FIGURE LEGENDS

**Supplemental Figure 1.** Changes in TM numbers during thymic development. Thymuses were harvested from mouse at various developmental time points and enzymatically digested to obtain single-cell suspensions. Cells were stained with antibodies for flow cytometry analysis. A) Total thymocyte numbers at different ages. B) Comparison of TM numbers and percentages across different ages of mice. Data are presented as mean ± SEM (n = 6-12 per group from 2-4 independent experiments).

**Supplemental Figure 2.** MaFIA transgene and reporter expression in thymic macrophages. A) The *Csf1r*-driven MaFIA transgene encodes EGFP together with a membrane-targeting low-affinity nerve growth factor receptor segment (dLNGFR) and an FKBP-Fas suicide module. dLNGFR is a structural component of the fusion construct and is not used as a lineage marker. B) AP20187 is a synthetic homodimerizer that binds the engineered FKBP domains, bringing Fas intracellular domains together and inducing apoptosis in transgene-expressing cells. C) Representative *Csf1r*-EGFP signal in MaFIA TMs (green) and C57BL/6J controls (gray). D-E) *Csf1r*-EGFP distributions and mean fluorescence intensity across TIMD4+ VCAM1+, TIMD4-VCAM1+, and TIMD4-VCAM1-populations. Data are mean ± SEM (n = 4 per group from two independent experiments), Kruskal-Wallis test (**p ≤ 0.01).

