## Supplementary figures and images for "Heterogeneity and Ontogeny of Mouse Thymic Macrophages Reveal a Requirement for *Csf1r*-Expressing Myeloid Cells During Early T Cell Development"

### Supplemental Figure

#### Supplemental Figure 1

**A**

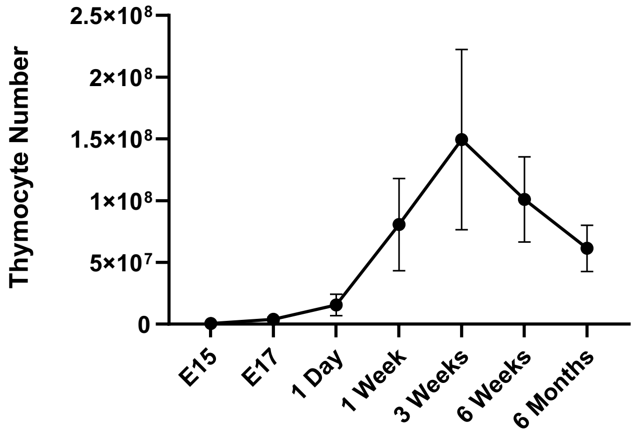


**B**

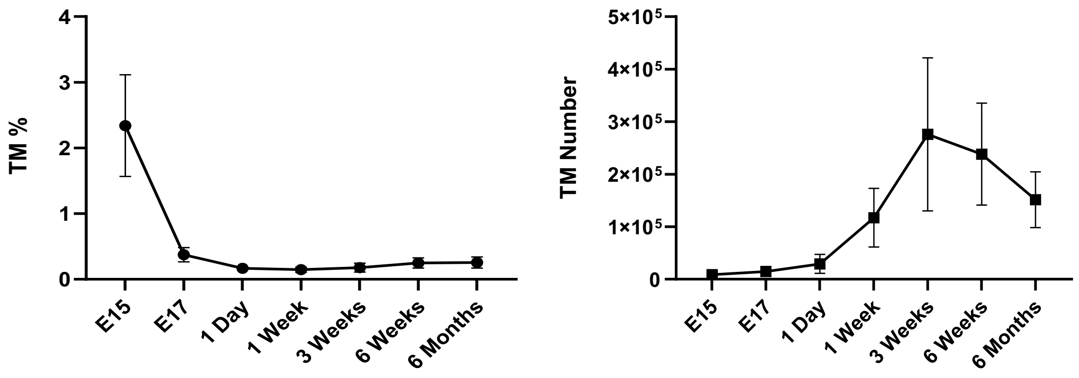

#### Supplemental Figure 2

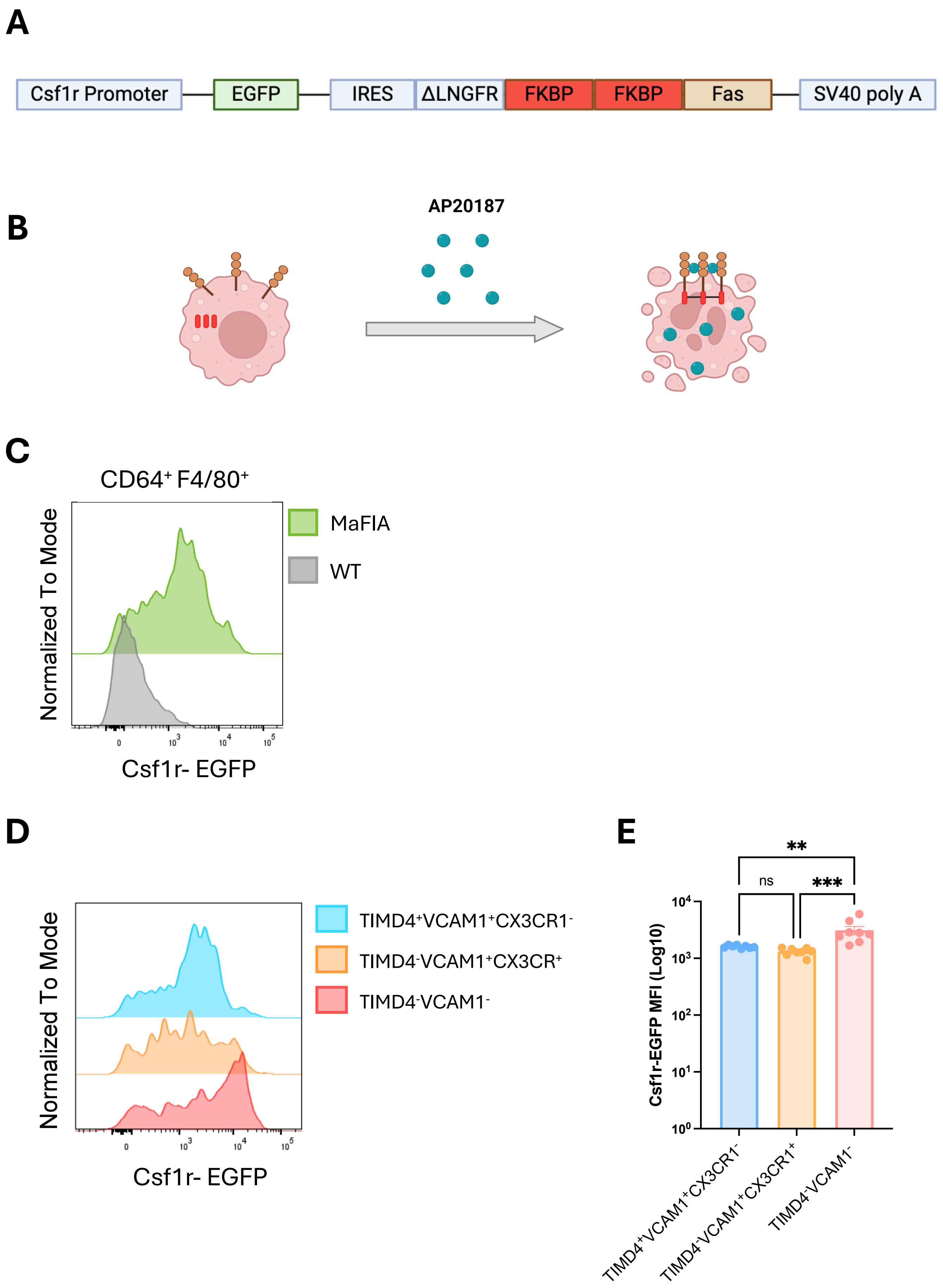
